# Impaired larval motor function in a zebrafish Stathmin-2 (STMN2) knockout model

**DOI:** 10.1101/2025.10.24.684380

**Authors:** Tyler J.N. Gurberg, Ziyaan A. Harji, Christian J. Rampal, Joséphine Sacy-Richer, Amy Wang, Esteban C. Rodríguez, Gary A.B. Armstrong

**Affiliations:** Department of Neurology and Neurosurgery, Montreal Neurological Institute, Faculty of Medicine, McGill University

## Abstract

Stathmin-2 (STMN2) is a microtubule associated protein that plays a role in the stability of microtubules of axons in the nervous system of animals. In this study we generated a novel zebrafish *STMN2* knockout (KO) model. *STMN2* is represented by two genes in the zebrafish genome: *stmn2a* and *stmn2b*. Using the CRISPR/Cas9 mutagenic system we selected founder fish lines harbouring frameshift mutations in both genes and bred these together to generate a double *stmn2a* and *stmn2b* KO model. Using these models, we observed increased developmental lethality in our double *stmn2a* and *stmn2b* KO model and impaired motor function at larval stages of development. Examination of the larval neuromuscular junction (NMJ) revealed a slight increase in the number of orphaned NMJs in trunk musculature as well as a reduction in amplitude of miniature endplate currents in our double *stmn2a* and *stmn2b* KO model. In a final series of experiments, we show impaired ventral root axon regrowth following transection in double *stmn2a* and *stmn2b* KO larvae. Our findings suggest that while not essential for motor axon development, loss of *stmn2a* and *stmn2b* expression perturbs the assembly of zebrafish NMJs during development resulting in a minor motor phenotype and impairs that ability to regenerate motor axons following injury.

## Introduction

During vertebrate development, differentiated spinal cord motor neurons extend axons to the periphery, where they form NMJs with muscle cells. For proper targeting of muscle cells to occur, growth cones and their axons to respond dynamically to extracellular cues by remodeling their microtubule networks. Microtubule networks undergo remodeling through phases of individual microtubule polymerization and depolymerization in a process known as dynamic instability (Mitchison and Kirschner, 1984). Microtubule associated proteins are a class of proteins involved with microtubule dynamics and include members of the Stathmin family of proteins, with humans possessing 4 members (STMN1-4). The Stathmin-family of proteins all possess a shared C-terminal Stathmin-like domain that bind tubulin in a phosphorylation dependent manner by JNK1 (Belmont and Mitchison, 1996; Tararuk et al., 2006). Unlike STMN1, STMN2-4 differ in that they possess an N-terminal motif responsible for vesicular membrane attachment motif and a targeting signal that directs them to the Golgi complex (Di Paolo et al., 1997; Chauvin and Sobel, 2015; Benarroch, 2021), and are expressed exclusively in the nervous system (Bièche et al., 2003).

The function of the Stathmin-family of proteins in motor axon development remains incompletely understood despite both human and mouse motor neurons abundantly expressing *STMN2* (Sun et al., 2015; Melamed et al., 2019) suggesting an important role for regulating the cytoskeleton of these cells. STMN2, (also known as SCG10) is upregulated during development and following axonal injury where it is anterogradely trafficked to growth cones (Stein et al., 1988; Shin et al., 2014). STMN2 acts as a microtubule-destabilizing protein by sequestering soluble tubulin dimers and inducing curvature at microtubule tips, promoting rapid depolymerization and shortening events known as catastrophes (Curmi et al., 1997; Gardner et al., 2013; Gupta et al., 2013). These events are essential for growth cone advancement and responsiveness to guidance cues (Grenningloh et al., 2004; Manna et al., 2007). Investigations examining individual microtubules *in vitro*, suggest that increasing the molar ratio of STMN2 to tubulin increases the frequency of catastrophe events and microtubule growth rates (Manna et al., 2007). In rat primary cultures of hippocampal cells, siRNA knockdown of *Stmn2* expression has been shown to impair neurite growth (Morii et al., 2006) and increase the growth cone surface area (Poulain and Sobel, 2010). Similarly, cultured dorsal root ganglion neurons from Stmn2 knockout (KO) mice exhibit impaired axon outgrowth (Krus et al., 2022). These KO mice also display elevated perinatal mortality, and survivors develop sensory and motor neuropathy by three months of age, including NMJ denervation (Krus et al., 2022).

To investigate the role of STMN2 in motor axon development we generated a zebrafish *stmn2* knockout model. The zebrafish genome contains two orthologs of *STMN2* (*stmn2a* & *stmn2b*). Using the CRISPR/cas9 system we generated a knockout models of each gene and bred our fish to generate a double KO model. Using these models, we investigated larval motor phenotypes. During zebrafish embryo development, motor axons rapidly grow and form NMJs in the axial trunk musculature used for swimming behaviour. Orchestration of this process must be particularly tuned to effectively establish the circuitry for larval motor escape responses. In this study we demonstrate that *stmn2a* & *stmn2b* double KO larvae display elevated developmental lethality, impaired motor function, a slight increase in the number of orphaned AchRs clusters in muscle cells and impaired motor axon regrowth following axotomy. Our findings suggest that while not essential for motor axon development, loss of *stmn2* expression perturbs the assembly of zebrafish NMJs during development.

## Methods

### Zebrafish housing and maintenance

Adult zebrafish (*Danio rerio*) of the Tübingen long fin (TL) were maintained according to standard protocols (Westerfield, 1995) at 28.5°C under a 14/10 hour light/dark cycle in the animal research facility of the Montreal Neurological Institute (MNI), at McGill University (Montreal, Quebec, Canada). All experiments were approved by MNI’s Animal Care Committee (ACC) and followed the guidelines set by the Canadian Council for Animal Care (CCAC). Animal care protocol approval # MNI-7890.

### Cas9 mRNA and guide RNA synthesis

Synthesis of *Cas9* mRNA and guide RNA (gRNA) was performed using previously described methods (Jao et al., 2013; Vejnar et al., 2016). Target gRNA sites were identified using CRISPRscan (Moreno-Mateos et al., 2015), synthesized using the T7 MEGAscript kit (Invitrogen), and purified by phenol-chloroform extraction and ethanol precipitation. A zebrafish codon-optimized Cas9 (pT3TS-nCas9n, Addgene plasmid # 46757) was linearized with XbaI overnight and 1 μg of linear template DNA was used for *in vitro* transcription of mRNA using the T3 mMESSAGE mMACHINE® (Invitrogen) kit and purified by phenol-chloroform extraction and ethanol precipitation. The following gRNA target sites were used, with the protospacer-adjacent motif (PAM) site underlined:

5’ GAAGAAGCTGGAGGCGGCTGAGG 3’ for *stmn2a*, located in exon 3.

5’ GAGGCACGCAAGAACATCATGGG 3’ for *stmn2b*, located in exon 2.

### CRISPR/Cas9 mutagenesis and screening for founder lines

*Stmn2a* KOs*, stmn2b* KOs, and double *stmn2a/b* KOs were generated using methods previously described by our laboratory (Armstrong et al., 2016). *Cas9* mRNA (100 ng/μL) and gRNA (100 ng/μL) were co-injected in ∼2 nL volumes into zebrafish embryos at the one-to-two-cell stage of development. DNA cutting efficiency was evaluated through High Resolution Melting (HRM) analysis of 24 one-to-two-day post-fertilization (1-2 dpf) embryos per gRNA target (Erali and Wittwer, 2010). The most efficient gRNA target sites were selected for the creation of individual zebrafish lines with disruption of *stmn2a* and *stmn2b* reading frames and raised as F0s. Adult F0s were outcrossed with wild type zebrafish and fertilized embryos were screened for indel transmission using HRM.

HRM primer sequences for verification of *stmn2a* line:

Forward: 5’ GGCCCACAGCATCACCTCT 3’

Reverse: 5’ CATTTGTCTGTGTTTGCACTAACCC 3’

Primer sequences for verification of *stmn2b* line:

Forward: 5’ CGCAGCATACAAAGAGAAGATGAAGG 3’

Reverse: 5’ TCCTTCTCCACTGTGTCTCATTACTGG 3’

### General screening and knockout line selection

Adult F1s zebrafish were individually separated and anesthetized with 1% tricaine (MS-222, Sigma) and a caudal fin clip was used to extract DNA. DNA was extracted (Extract-N-Amp™ Tissue PCR Kit, Millipore Sigma), and target sites were amplified by polymerase chain reaction (PCR) and followed by restriction fragment-length polymorphism (BbvCI and NlaIII for *stmn2a* and *stmn2b*, respectively). Sequencing (Genome Québec) was performed on amplicons to identify the specific mutation of each founder line. Founder zebrafish carrying frameshift mutations resulting in premature stop codons were selected and homozygous carriers of each mutation were generated by in-crossing F2s to generate our *stmn2a* KO, *stmn2b* KO, and double *stmn2a/b* KO lines. Transgenic Tg[*Hb9:GFP*] fish, expressing GFP in their motor neurons, were crossed into the KO models to generate a double *stmn2a^-/-^*; *stmn2b^-/-^*; Tg[*Hb9:GFP*] line.

Primer sequences for verification of *stmn2a* line:

Forward: 5’ TAAAAGGGCCTCAGGTCAGG 3’

Reverse: 5’ GTTCTCAATGTGGCTGTCTTTTCC 3’

Primer sequences for verification of *stmn2b* line:

Forward: 5’ GTACCCTCTGGCTTAAAACGTAGATG 3’

Reverse: 5’ CCTCTCACTCATGAAACGAGCTTG 3’

### Larval motor function

Larval motor function was assessed using methods previously described (Armstrong and Drapeau, 2013a). Individual larvae aged 52-54 hours post-fertilization (hpf) were placed in the center of an aquatic arena (150 mm petri dish) containing fresh system water, temperature controlled between 24 °C and 25 °C. Individual larvae were allowed to habituate to the environment for 30-60 seconds. A light touch to the tail of the larvae was applied using a pair of forceps, evoking a burst swimming motor response. Movement was recorded from above at 30 Hz (Grasshopper 2 camera, Point Gray Research) until larvae either stopped moving or until they reached the side of the arena. Total swim distance and mean swim velocities were quantified following manual movement tracking (Manual Tracking plugin, ImageJ).

### RNA extraction and quantitative real time PCR (qRT-PCR)

RNA was extracted from 30 pooled 2 dpf larvae using the trizol-chloroform method. Purity and quality of RNA was verified using a NanoDrop® instrument. 2.5 μg of RNA was used to create a cDNA library using the SuperScript VILO cDNA Synthesis Kit (Invitrogen). Quantitative real-time PCR was performed on the cDNA libraries using the SYBR Green Supermix (Bio-Rad Laboratories). The delta-delta Ct method of relative quantification was performed with *gapdh* as an internal control (Livak and Schmittgen, 2001).

Primer sequences for RT-qPCR analysis of *stmn2a* line:

Forward: 5’ GAACAAGGAAAACCGTGAGG 3’

Reverse: 5’ CTTGTTCCTGCGGACGAT3’

Primer sequences for RT-qPCR analysis of *stmn2b* line:

Forward: 5’ CTGAAAGCCATGGAGGAGAA 3’

Reverse: 5’ ACCAGCTGAGCGTGCTTC 3’

### Neuromuscular junction colocalization and axonal microtubule networks

Immunofluorescent microscopy experiments were performed to assess NMJ integrity. Double labelling of pre-synaptic and post-synaptic membranes was achieved using a synaptotagmin 2 (Syt2, pre-synaptic) antibody and sulforhodamine-conjugated alpha-bungarotoxin markers (α-Btx, post-synaptic), as previously described (Armstrong and Drapeau, 2013a). Briefly, 10 dechorionated 2 dpf zebrafish larvae were fixed in 4% paraformaldehyde (PFA) in phosphate-buffered saline solution (PBS) overnight at 4 °C with a gentle rotation. Larvae were then rinsed with PBS (3 x 15 minutes) and incubated with 1 mg/mL collagenase in PBS solution for 45 minutes at room temperature (RT). Larvae were again rinsed in PBS (3 x 15 minutes) and incubated in PBST (PBS, Triton X-100) for 30 minutes at RT and then incubated in a 10 mg/mL sulforhodamine-conjugated α-Btx in PBST solution for 30 minutes at RT. Following this, larvae were rinsed with PBST (3 x 15 minutes) and incubated for an hour in fresh blocking solution (2% Goat serum, 1% bovine serum albumin (BSA), 0.1% Triton X-100, 1% DMSO in PBS) at RT before being left overnight in block solution containing the Syt2 primary antibody (DSHB, 1:100). The next day larvae were rinsed with PBST (4 x 15 minutes) and incubated at RT for 5 hours in fresh blocking solution containing the secondary antibody (Alexa fluor 647, ThermoFisher, 1:1200). Larvae were rinsed with PBST (3 x 15 minutes) and incubated in PBST overnight at 4 °C with a gentle rotation and then placed in a solution containing 70% glycerol and mounted on glass slides for imaging. NMJs were visualized with a 60x/1.42 oil immersion objective on a Quorum Technologies microscope with an 89 NORTH LDI spinning disk confocal mounted on an Olympus BX61W1 fluorescence Microscope, connected to a photometrics prime BSI camera. Images were acquired using Volocity software (Improvision).

### Electrophysiology

Using previously described methods (Buss and Drapeau, 2002), zebrafish were anaesthetized in 0.04% tricaine (Sigma) dissolved in modified Evans solution containing, in mM: 134 NaCl, 2.9 KCl, 2.1 CaCl2, 1.2 MgCl2, 10 HEPES, 10 glucose, adjusted to 290 mOsm and pH 7.8. Larvae were then pinned using fine (0.001 in.) tungsten wires through their notochords to a Sylgard-lined dish. The outer layer of skin between the pins was removed using a fine glass electrode and forceps exposing the trunk musculature. The preparation was then visualized by oblique illumination (Zeiss Examiner A1 upright microsope) and standard whole-cell voltage clamp recordings were obtained from fast twitch (embryonic white) muscle cells identified visually by their oblique orientation to the body axis. 5-6 MΩ glass electrodes were pulled from thin-walled Kimax-51 borosilicate glass (Kimble Chase) and filled with the following intracellular solution (in mM: 130 CsCl, 2 MgCl2, 10 HEPES, and 10 EGTA adjusted to pH 7.2, 290 mOsm.). Cells were held near their resting potential at -65mV and series resistant was < 8 MΩ and compensated to 70-90 %. 1 µM TTX was perfused over the preparation to isolate spontaneous (quantal) miniature endplate currents (mEPC) over a 10 minute recording period. All electrophysiological data were sampled at 40 kHz using an Axopatch 200B amplifier (Molecular Devices) and digitized using a Digidata 1550B (Molecular Devices) and later analyzed using pCLAMP 10 software (Molecular Devices).

### Axon transection and recovery

Wild type and double *stmn2a*^-/-^; *stmn2b*^-/-^ larvae expressing a single copy of the Tg[*Hb9:GFP*] transgene was selected for axon ablation. Selected larvae were embedded in 1% low melting point agarose, prepared using system water, at 50 hpf. Pre-transection images of fluorescently labelled ventral root axon projections were collected. A microinjection needle was then used to fully transect an individual ventral root axonal projection in a single hemisomite between somites 13-17. Cut ventral roots were re-imaged every hour for 24 hours using a 60x/1.42 water immersion objective.

### Statistical analyses

Statistical analyses were performed using Prism 9 (GraphPad Software Inc.). Shapiro-Wilks test was used to test for normal sample distributions. Tests with only two samples were compared using unpaired students’ t-tests. Kruskal-Wallis tests were used to compare data sets containing more than two non-normally distributed samples, followed by Dunn’s post-hoc multiple comparisons test. Chi-squared tests were used to compare genotypic frequencies. Significance was assessed at *p* < 0.05.

## Results

### Generation of stmn2a, stmn2b, and double knockout zebrafish lines

The gene encoding STMN2 is represented by two orthologs in the zebrafish genome: *stmn2a* and *stmn2b* (**Fig. 1A**). Guide RNAs targeting the coding regions within exons 2 and 3 for each gene were individually evaluated for cutting efficiency using the CRISPR/Cas9 mutagenic system. HRM analysis of F1 offspring revealed that a gRNA targeting exon 3 of *stmn2a,* and a gRNA targeting exon 2 of *stmn2b,* were robustly cutting DNA at these loci and where subsequently selected for generating our mutant fish lines. Sanger sequencing of F2 homozygous larvae revealed an 11-nucleotide deletion in exon 3 of *stmn2a* and a 13-nucleotide deletion in exon 2 of *stmn2b* (**Fig. 1B**), resulting in premature stop codons in the open reading frame of their respective transcripts that could be screened by the loss of restriction enzyme sites in amplified PCR products (**Fig. 1C**). A double *stmn2a* and *stmn2b* KO fish line was generated by inbreeding both lines. Additionally, a double KO fish line expressing the Tg[*Hb9:GFP*] transgene (Flanagan-Steet et al., 2005) was also crossed into our mutant lines to create a double KO & Tg[*Hb9:GFP*] transgenic fish line. To confirm that the mutations were disrupting gene function we quantified *stmn2a* and *stmn2b* transcript levels using cDNA extracted from RNA collected in from pooled 2 dpf larvae. Both homozygous mutant lines showed significantly reduced expression of *stmn2a* and *stmn2b* transcripts suggesting that the transcripts were likely removed through nonsense-mediated decay (**1D**). As both fish Stmn2a and Stmn2b are similar to one another (**Fig 2**), we speculated that each paralog might be able to compensate for the loss of expression of the other by displaying elevated transcriptional expression. However, examination of transcript levels from whole 2 dpf larvae, revealed that there was no change in the expression of *stmn2a* in *stmn2b*^-/-^ zebrafish and a trend of increased expression of *stmn2b* in *stmn2a^-/-^* cDNA samples but this did not reach significance (**Fig. 1D**). Examination of 2 dpf larval genotype frequency following a double heterozygous incross (*e.g., stmn2a*^+/-^ ; *stmn2b*^+/-^) revealed that fewer double KO larvae (*e.g., stmn2a*^-/-^ ; *stmn2b*^-/-^) were surviving to this stage of development (**Fig 1E**). We next investigated if there were changes in larval survival over the first 30 days of development. We observed that double KO larvae displayed impaired survival when compared to wild type larvae during the first month of development (**Fig 1F**). Surviving double KO zebrafish were able to reach sexual maturity normally, however adult male double *stmn2a*^-/-^ ; *stmn2b*^-/-^ zebrafish were behaviourally unable to fertilize embryos of any genotype (likely due to poor adult motor function) but could be used for *in vitro* fertilization to generate clutches of double *stmn2a*^-/-^ ; *stmn2b*^-/-^ embryos.

**Figure 1.**
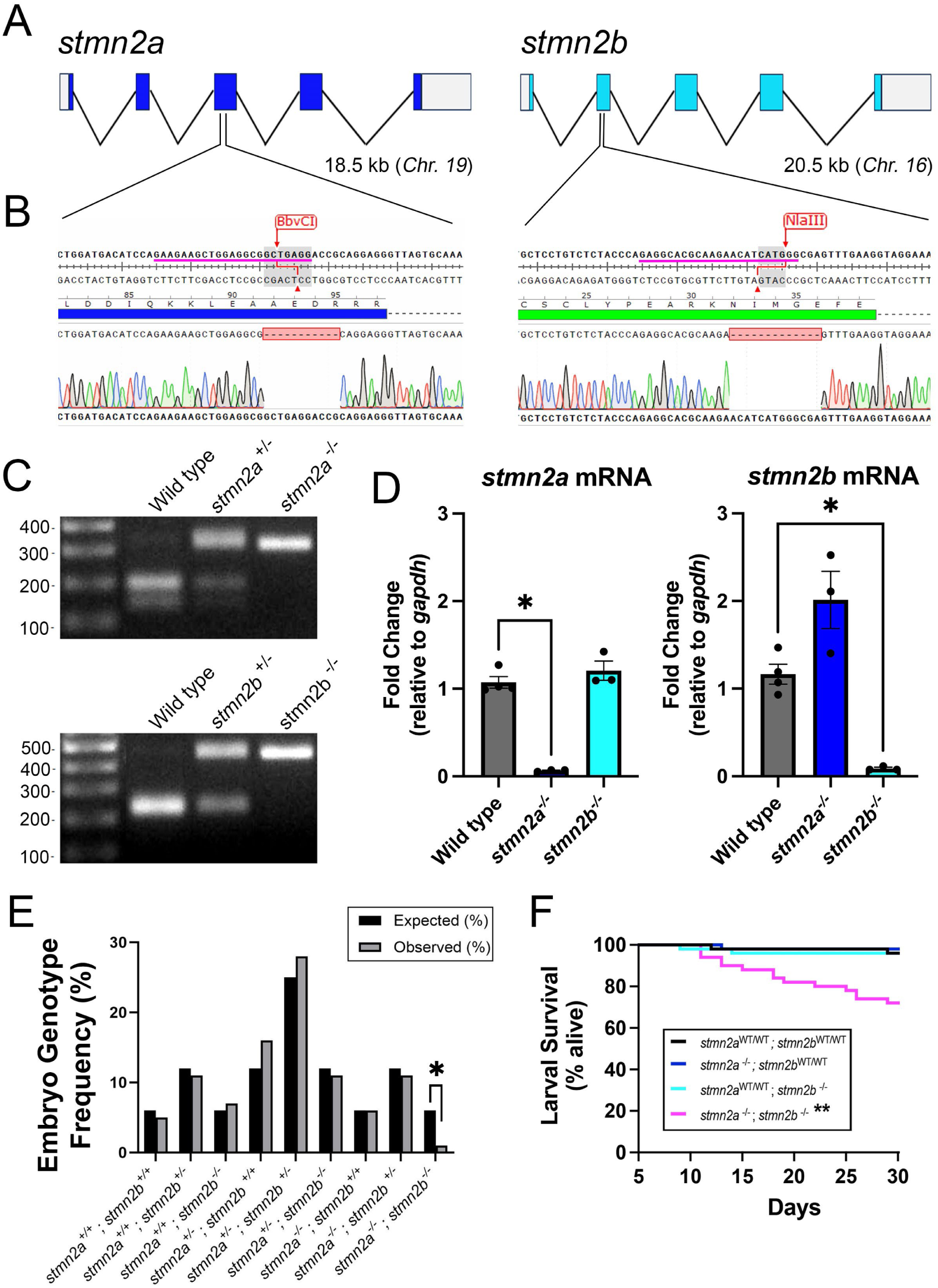
Development of the *stmn2a* and *stmn2b* KO zebrafish lines. **A**, Schematic representation of zebrafish *stmn2a* and *stmn2b* genes showing gene structure. **B**, Location of guide RNAs (pink lines) for each gene and mutant lines selected for use. An 11-nucleotide deletion and a 13-nucleotide deletion were produced in *stmn2a* and *stmn2b*, respectively, resulting in premature stop codons in the open reading frame. **C**, Loss of BbvCI (*stmn2a*) and NlaIII (*stmn2b*) restriction enzyme digestions sites at their respective gRNA target sites were used for genotyping of digested amplicons. **D**, Analysis of gene expression using RT-qPCR of 2 dpf larvae, relative to *gapdh* expression. A significant reduction in *stmn2a* expression in *stmn2a^-/-^* larvae, and a significant reduction in *stmn2b* in *stmn2b^-/-^* larvae was observed, suggesting the mutant transcripts are targeted for nonsense-mediated decay. No *stmn2a* compensation was observed in the *stmn2b^-/-^* larvae, whereas a trend towards *stmn2b* compensation in *stmn2a^-/-^*larvae was observed. Statistical analysis by Mann-Whitney test, significance determined at *p* < 0.05. Sample sizes; wild type, N = 4 batches of 30 pooled larvae; *stmn2a^-/-^,* N = 3; *stmn2b^-/-^,* N = 3. **E**, A significant reduction in the expected frequency of double *stmn2a*^-/-^; *stmn2b*^-/-^ 2 dpf larval was identified following a *stmn2a*^+/-^; *stmn2b*^+/-^ X *stmn2a*^-/-^; *stmn2b*^+/-^ incross when compared to the expected frequency (*p* < 0.05). **F**, Survival curve showing a reduction in *stmn2a*^-/-^; *stmn2b*^-/-^ larval survival over the first 30 days of development (*p <* 0.01). * Represents (*p* < 0.05), ** (*p* < 0.01).

**Figure 2.**
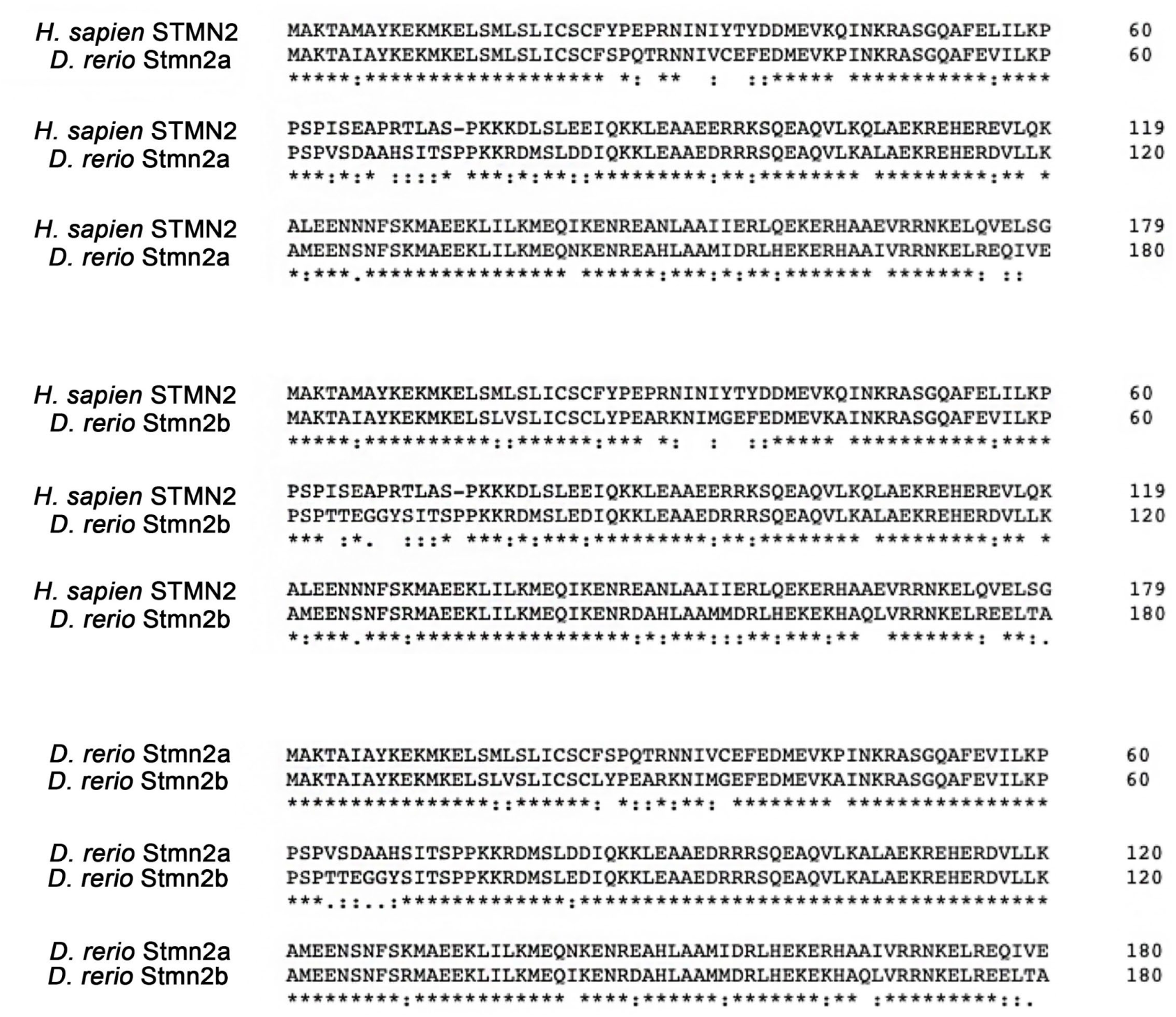
Amino acid sequence alignment between human STMN2 and zebrafish Stmn2a (top), human STMN2 and zebrafish Stmn2b (middle), and zebrafish Stmn2a and Stmn2b (bottom). Zebrafish Stmn2a shares 78% identity and 90% similarity with human STMN2, while Stmn2b shares 73% identity and 88% similarity with human STMN2. Zebrafish Stmn2a and Stmn2b and are 85% identical and 93% similar to each other.

### Stmn2 KO zebrafish display a significant reduction in larval motor function

Given that both *stmn2a* and *stmn2b* transcripts are expressed during development, we next sought to determine whether the loss of expression of either transcript affected larval motor function. At 2 dpf, zebrafish larvae display burst swimming behaviour in response to a light tail touch (Buss and Drapeau, 2001). Using zebrafish aged 50-54 hpf, we conducted touch-response assays in an aquatic arena (**Fig 3 A, B**), following previously established methods (Armstrong and Drapeau, 2013a; Armstrong and Drapeau, 2013b). Larvae in all genetic groups were able to respond to the light touch to the tail. Examination of motor parameters revealed a significant reduction in mean swim velocity in *stmn2a*^-/-^, *stmn2b*^-/-^, and double *stmn2a*^-/-^; *stmn2b*^-/-^ larvae when compared to controls (**Fig. 3C**). Furthermore, total swim distance was significantly reduced in *stmn2a*^-/-^ and double *stmn2a*^-/-^; *stmn2b*^-/-^ larvae when compared to wild type larvae. Whereas *stmn2b*^-/-^ larvae exhibited swim distances that were that were similar to wild type controls (**Fig 3D**).

**Figure 3.**
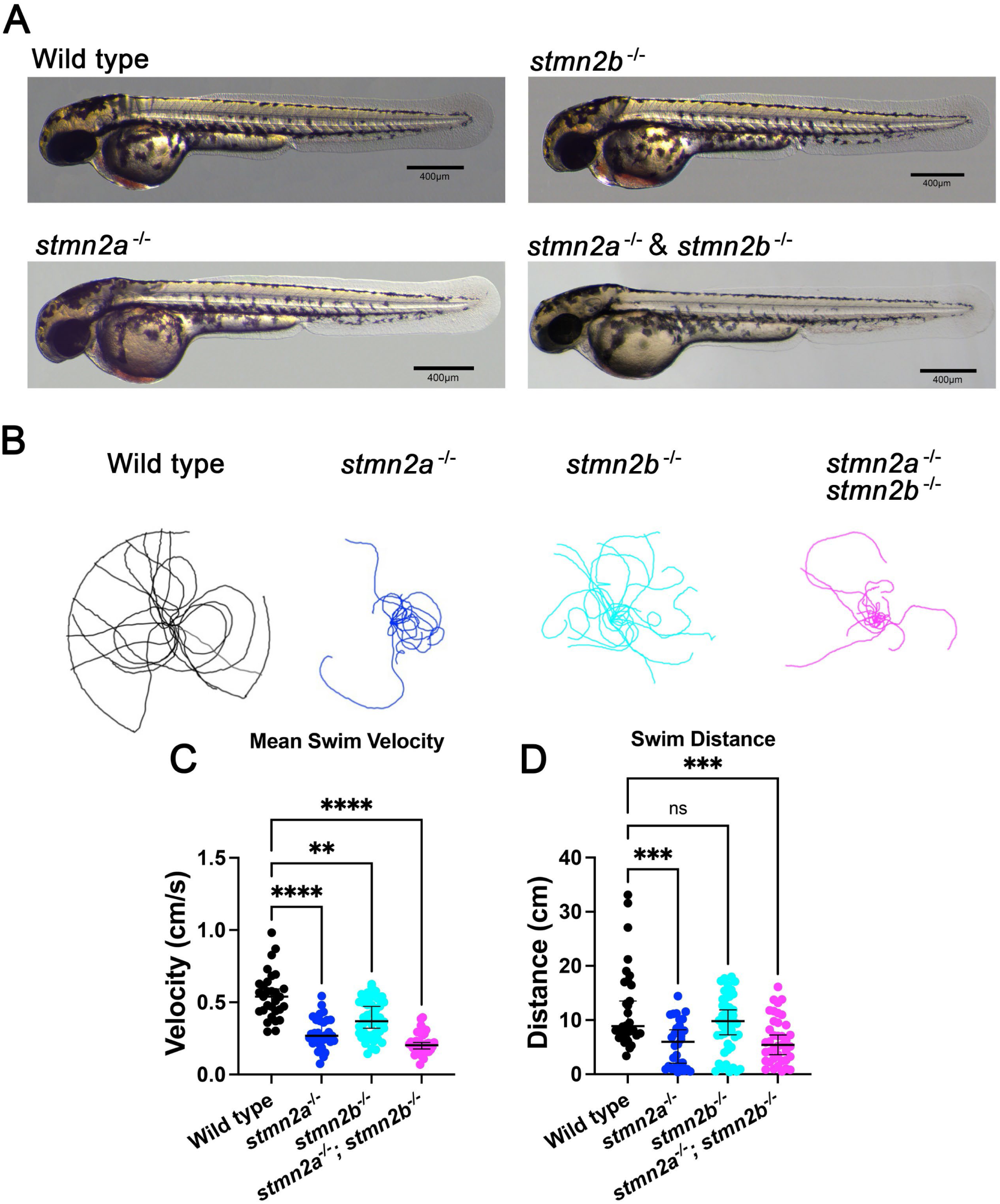
Two-day old larval touch-evoked motor response assays for *stmn2a*^-/-^, *stmn2b*^-/-^, and double *stmn2a* ^-/-^; *stmn2b*^-/-^ zebrafish. **A**, Example images of 2 dpf larvae. Scale bar represents 400 µm. **B**, Fifteen individual superimposed larval motor traces from wild type, *stmn2a*^-/-^, *stmn2b*^-/-^, and double *stmn2a*^-/-^ ; *stmn2b*^-/-^ genetic groups. **C**, Tabulation of mean swim velocity and swim distance. **D**, Mean swim velocity ± standard error and sample sizes were as follows: wild type (0.55 ± 0.03 cm/s, n = 30); *stmn2a*^-/-^ (0.9 ± 0.02 cm/s, n = 30); *stmn2b*^-/-^ (0.39 ± 0.02 cm/s, n = 45); and double *stmn2a*^-/-^ & *stmn2b*^-/-^ (0.22 ± 0.01 cm/s, n = 39). (**C**), Mean swim distances, standard error, and sample sizes were as follows: wild type (12.4 ± 1.4 cm, n = 30); *stmn2a*^-/-^ (5.7 ± 0.8 cm, n = 30); *stmn2b*^-/-^ (9.3 ± 0.8 cm, n = 45); double *stmn2a*^-/-^ & *stmn2b*^-/-^ (6.3 ± 0.7 cm, n = 39). Kruskal-Wallis multiple comparisons tests were used to determine significance. * Represents (*p* < 0.05), ** (*p* < 0.01), and *** (*p* < 0.005) and **** (*p* < 0.001).

### Double stmn2a^-/-^; stmn2b^-/-^ zebrafish present decreased colocalization of pre- and post- synaptic NMJ markers

As motor function impairment could result from abnormal NMJs, we next investigated if there were defects at these sites. Emanating from the larval spinal cord, ventral roots in 2 day old zebrafish are composed of 20-30 axons from primary and secondary spinal motor neurons (Myers, 1985; Westerfield et al., 1986) that project to and innervated axial muscles in each hemisegment. Reduced NMJ integrity has been described in previous ALS-mutant zebrafish models, where receptors can be considered ‘orphaned’ when post-synaptic puncta (*e.g*., nicotinic acetylcholine receptors, AchR) do not co-localize with pre-synaptic motor neuron markers (*e.g.,* synaptotagmin 2, Syt2) (Armstrong & Drapeau, 2013; Bose et al., 2019). To investigate larval trunk NMJs in our mutant *stmn2* models we performed immunofluorescence confocal imaging of fixed zebrafish (aged 55-56 hpf) and labelled pre- and post-synaptic components with an antibody targeting Syt2 and alpha-bungarotoxin conjugated to sulforhodamine (aBtx) targeting AchRs (**Fig 4A**). We observed a significant increase in the number of orphaned pre-synaptic Syt2 puncta in *stmn2a*^-/-^ and double *stmn2a*^-/-^; *stmn2b*^-/-^ zebrafish, suggesting that the motor axons in these larvae possessed hyperbranched axon terminals that did not project to AchRs clusters (**Fig 4B**). Conversely, the number of orphaned post-synaptic AchR puncta generally did not differ among our models but we did observe a slight increase in the number of orphaned AchR puncta in our double *stmn2a*^-/-^; *stmn2b*^-/-^ fish model when compared to wild type NMJs (**Fig. 4C**). To further investigate the nature of NMJ functionality we recorded miniature endplate currents (mEPCs) on fast-twitch muscle cells of the zebrafish trunk (somites 10-15). In these recordings we observed that mEPCs amplitude in double *stmn2a*^-/-^; *stmn2b*^-/-^ larvae was reduced in comparison to recordings from aged-matched wild type larvae (**Figure 4D, E**) with no change being observed in the frequency of mEPCs (**Figure 4F**). Additionally, we observed no differences between wild type and double *stmn2a*^-/-^; *stmn2b*^-/-^ larvae in our measures of mean 10-90% rise time (1.35 ± 0.19 *vs* 1.75 ± 0.09 ms), mean decay (τ) constant (13.52 ± 1.44 *vs* 12.99 ± 2.30), mean resting muscle membrane potential (-62.75 ± 2.71 *vs* -65.83 ± 2.51), or mean muscle membrane capacitance (69.38 ± 12.34 *vs* 76.02 ± 8.05 pF).

**Figure 4.**
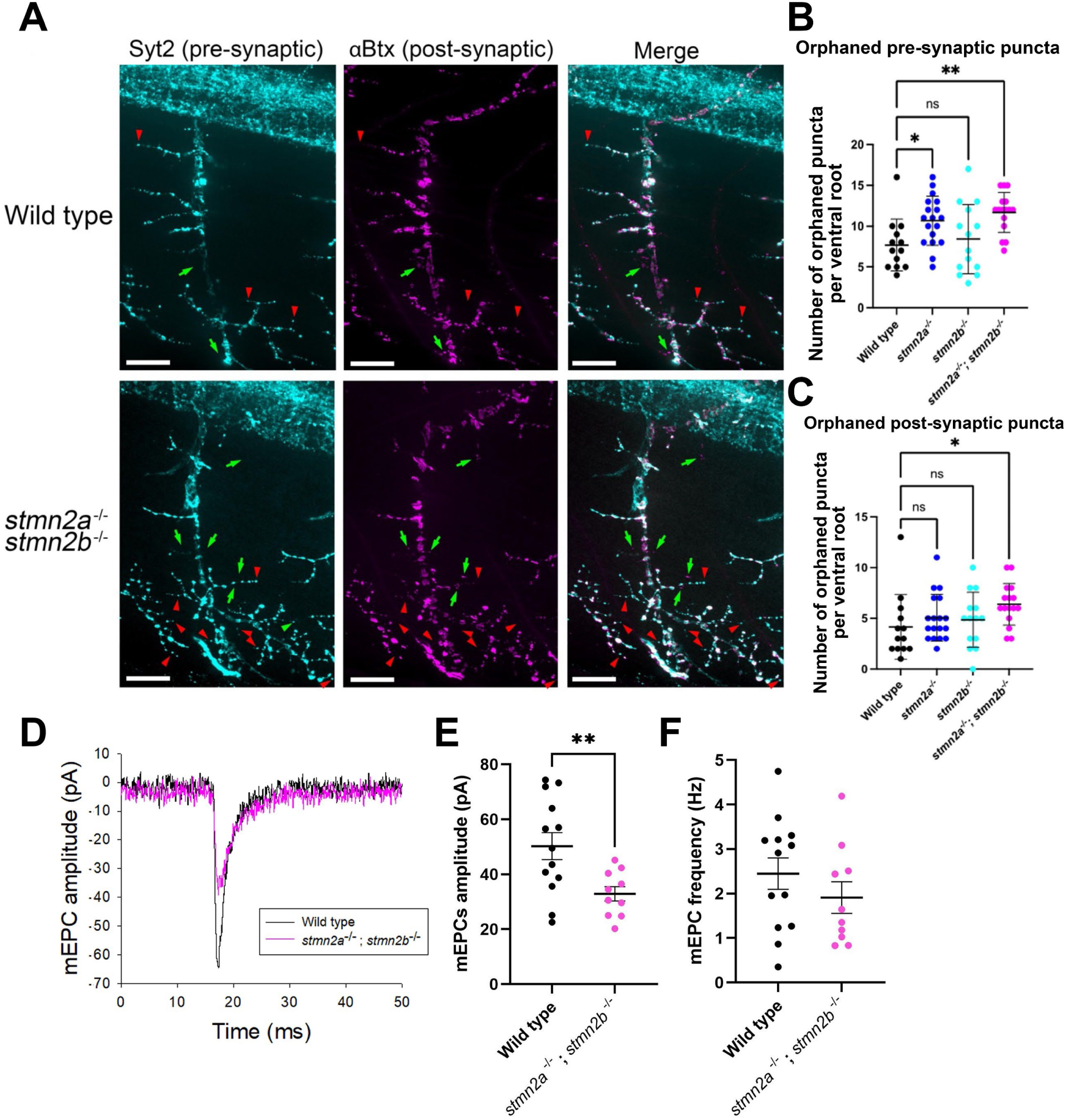
Two-day old larval zebrafish NMJs from double *stmn2^-/-^*and *stmn2b^-/-^* zebrafish display increased pre- and post-synaptic puncta and decreased quantal size of mEPCs (e.g., mEPC amplitude). **A**, Representative immunofluorescent images of larval NMJs from wild type and double *stmn2a^-/-^* and *stmn2b^-/-^*larvae. Scale bars represent 25 µm. Quantification of orphaned pre-synaptic Syt2 puncta (**B**) and post-synaptic AchR puncta (**C**). Mean number of orphaned pre-synaptic terminals ± standard error and sample sizes were as follows: wild type (7.7 ± 0.9 terminals, n = 13); *stmn2a^-/-^* (10.7 ± 0.7 terminals, n = 18); *stmn2b^-/-^* (8.4 ± 1.1 terminals, n = 14); double *stmn2a^-/-^* & *stmn2b^-/-^* (11.7 ± 0.6 terminals, n = 16). Mean number of orphaned post-synaptic terminals ± standard error, and sample sizes were as follows: wild type (4.2 ± 0.9 terminals, n = 13); *stmn2a^-/-^* (5.1 ± 0.5 terminals, n = 18); *stmn2b^-/-^* (4.9 ± 0.7 terminals, n = 14); double *stmn2a^-/-^* & *stmn2b^-/-^* (6.4 ± 0.5 terminals, n = 16). **D**, Example mEPC events in wild type and double *stmn2a^-/-^* and *stmn2b^-/-^*larvae. **E**, Quantification of mean mEPC amplitude. **F**, Quantification of mean mEPC frequency. Samples sizes for mEPC recordings were as follows: wild type = 13; double *stmn2a^-/-^* & *stmn2b^-/-^* = 10. Data points represent individual means from larval recordings with black bar representing the overall group mean ± standard error. A Kruskal-Wallis multiple comparisons tests were used to determine significance B & C) or a t-test (E & F). * Represents (*p* < 0.05), ** (*p* < 0.01).

### Double stmn2a^-/-^; stmn2b^-/-^ zebrafish display impaired nerve regeneration following ventral root nerve injury

As ventral root projections in our double *stmn2a* and *stmn2b* KO larval model only displayed minor NMJ defects, we next explored if there were abnormalities in axon regeneration following injury. Using restrained larvae, aged 50 hpf, we transected single ventral root projections using a fine glass microelectrode and then recorded axon regeneration over a 24 hr period. Ventral root axon projections were visualized using larvae expressing the Tg[*Hb9:GFP*] transgene (**Fig. 5A**). Wild type zebrafish expressing the Tg[*Hb9:GFP*] transgene displayed fully degenerated distal sections of the ventral root projection within 3-4 hrs post injury (hpi) (**Fig. 5A**). However, in double *stmn2a^-/-^* ; *stmn2b^-/-^* larvae expressing the Tg[*Hb9:GFP*] transgene (**Fig. 5B**) transected ventral root project took longer to fully degenerate (6-12 hpi). Examination of the ventral root projection at 24 hpi revealed that 7/8 wild type motor neurons had fully regrown their axon projections whereas none of our double *stmn2a^-/-^* ; *stmn2b^-/-^* larvae were able to regrow their ventral root axon projections. These findings suggest a significant role for Stmn2a/b in motor axon maintenance and regeneration in zebrafish larvae.

**Figure 5.**
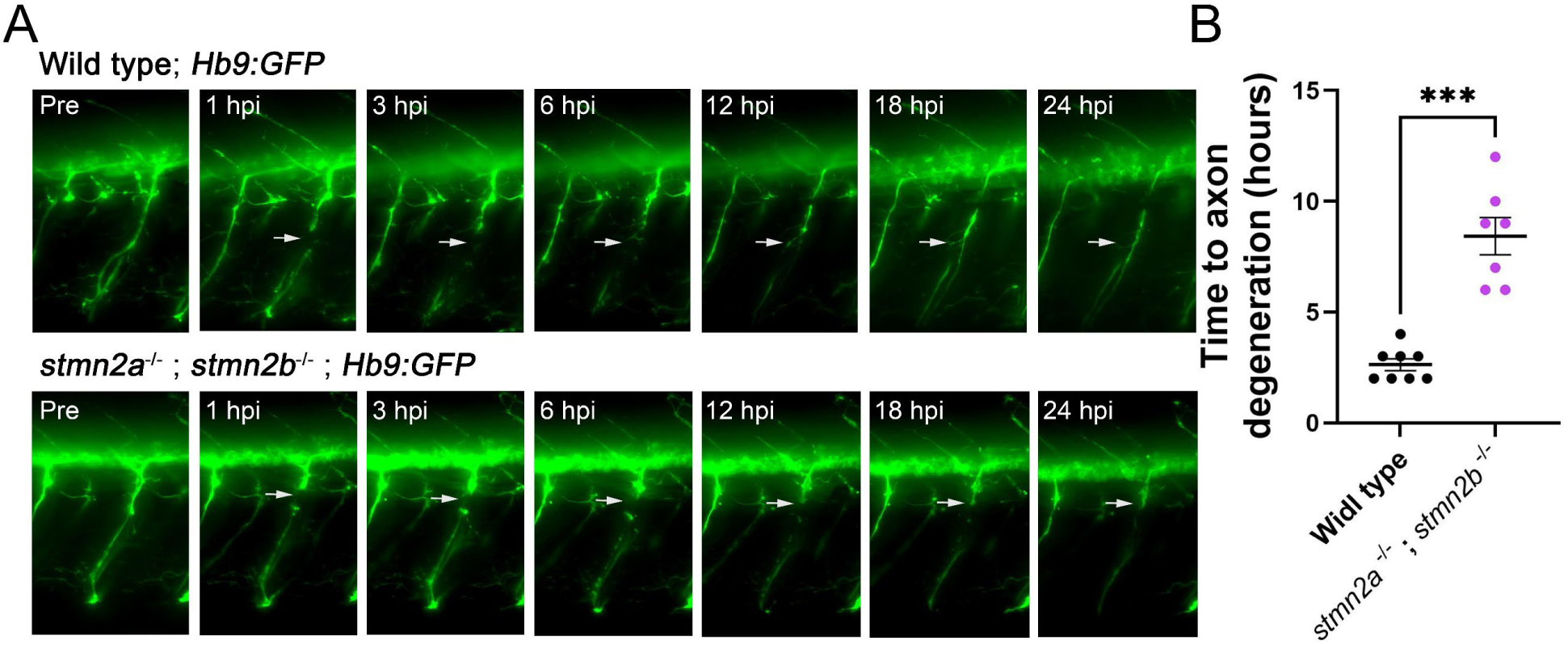
Delayed ventral root motor axon degeneration following axotomy in 2 dpf *stmn2a*^-/-^ ; *stmn2b*^-/-^ zebrafish. **A**, Example confocal images of larval ventral root motor axons of a wild type larvae (top) expressing Tg[*Hb9:GFP*] and *stmn2a^-/-^* ; *stmn2b^-/-^* larvae (bottom) expressing Tg[*Hb9:GFP*] over a 24 hour period following injury (hpi). White arrows indicate location of axotomy. **B**, Quantification of the length of time it took for distal motor axon degeneration to occur. Axon degeneration time was significantly longer in *stmn2a^-/-^* ; *stmn2b^-/-^* larvae when compared to wild type zebrafish. Sample sizes are as follows: wild type, n = 9; *stmn2a^-/-^* ; *stmn2b^-/-^*, n = 8). Data points represent individual axon degeneration times from different larvae with black bar representing the overall group mean ± standard error. A *t*-test was used to make statistical comparisons. *** Represents (*p* < 0.001).

## Discussion

Using CRISPR/Cas9 mutagenesis we developed a zebrafish *stmn2* knockout model and investigated abnormalities in the motor system at larval stages of zebrafish development. To generate our KO model, we selected mutant zebrafish lines containing an 11-nucleotide deletion in exon 3 of *stmn2a*, and a 13-nucleotide deletion in exon 2 of *stmn2b*, both of which produce premature stop codons that resulted in their mRNA transcripts being targeted by nonsense-mediated decay (**Fig 1D**). Single-cell RNA-Seq data (Sur et al., 2023) and *in situ* hybridization experiments (Burzynski et al., 2009) suggest increased *stmn2a* expression relative to *stmn2b* in the developing zebrafish. To examine if either *stmn2a* or *stmn2b* could compensate for loss of the other, we examined levels of their transcripts in our single gene KO lines. We found no changes in the levels of *stmn2a* in our *stmn2b*^-/-^ line, however, there was a trend of increased *stmn2b* transcript levels in our *stmn2a*^-/-^ line. This might suggest that *stmn2a* expression plays a more prominent role during development. While in mammalian *Stmn2* KO models compensation from other stathmin-family proteins has not been observed, it is unclear if a similar finding would be observed in zebrafish larvae. However, one line of evidence that compensation from other Stathmin proteins was not occurring in our *stmn2* KO model comes from our observations of reduced survival in double *stmn2a^-/-^* ; *stmn2b^-/-^* zebrafish during developmental (**Fig 1E, F**). Despite this early survival defect in our double *stmn2a^-/-^* ; *stmn2b^-/-^* model we were able to raise some fish to adulthood however we observed that males were behaviourally unable to fertilize female zebrafish of any genotype. *In vitro* fertilization using double *stmn2a^-/-^* ; *stmn2b^-/-^* sperm confirmed the male gametes are able to fertilize eggs, however their consistent inability to breed naturally is noteworthy and likely reflects impaired motor function.

As both orthologs of *stmn2* are expressed during development and are thought to play a role in microtubule stability in motor axons, we next investigated if there were abnormalities in larval motor function (**Fig 3**). While all our models were able to respond to a light touch to the tail evoking a motor touch response, we did observe that *stmn2a^-/-^*, *stmn2b^-/-^* and double *stmn2a^-/-^*; *stmn2b^-/-^* larval displayed a reduction in our measure of mean swim velocity. Whereas only our *stmn2a^-/-^*and double *stmn2a^-/-^; stmn2b^-/-^* models displayed a reduction in total swim distance. These findings suggest that while not essential for motor function, loss of *Stmn2* expression is coincident with a slight impairment in motor function at larval stages of zebrafish development. To investigate potential causes of impaired motor function we next investigated if there were abnormalities at larval NMJs. Examination of pre-synaptic and post-synaptic receptor NMJ clusters in the larval trunk musculature revealed that double *stmn2a^-/^*^-^ ; *stmn2b^-/-^* larvae possessed more pre- and post-synaptic puncta labelling (**Fig 4**). However, the number of pre- and post-synaptic NMJs, that did not colocalization with each other, was increased in our double *stmn2a^-/-^*; *stmn2b^-/-^*model. Our single gene KO models displayed NMJs that were similar to wild type with one exception being that *stmn2a^-/-^* larval trunks displayed an increase in the number of orphaned pre-synaptic puncta. To further investigate NMJ function we recorded mEPCs from fast-twitch muscle cells. We observed a reduction in the amplitude of mEPCs in double *stmn2a^-/-^*; *stmn2b^-/-^*larvae and no change in any other metric of mEPCs when compared to wild type larvae recordings (**Fig 4D-F**). This suggest that while NMJs are able to be formed in our model, the synaptic strength of individual NMJs is reduced. An alternative possibility is there is a defect in the clustering of AchRs on muscle cells in our double *stmn2a^-/-^* ; *stmn2b^-/-^* model. Muscle cell abnormalities, including the presence of increased centrally located nuclei suggesting muscle cell regeneration and fragmented NMJs, have been reported in *Stmn2* KO mice (Guerra San Juan et al., 2022). As both *stmn2a* and *stmn2b* are widely expressed in the larval CNS it maybe that other cell types, in particular neurons that process sensory information are impaired. However, none of our larvae failed to react to the tail touch, suggesting that at this stage in development the sensory system of the larvae is intact. This contrasts to what have been reported in multiple mouse *Stmn2* KO models (Krus et al., 2022; Li et al., 2023; Lopez-Erauskin et al., 2023). At this stage in zebrafish development, mechanosensation is processed by spinal cord Rohon–Beard cells and only later in development is sensory processing replaced by cells of the DRG (Kuwada et al., 1990) (Ribera and Nüsslein-Volhard, 1998). As such it maybe that loss of Stmn2 expression does not affect Rohon-Bear cells. While we did not examine sensory function at later stages in development it maybe that defects arise in older fish when cells of the DRG take over sensory processing.

In a final series of experiments we investigated if there were defects in motor axon regeneration following injury in our double *stmn2a^-/-^* ; *stmn2b^-/-^* larval model. Following motor axon transection, we observed a slower time course for distal regions of the motor axon to degenerate (**Fig 5**). This contrast findings to a study examining mouse dorsal root ganglion (DRG) axon degeneration that suggested faster axon degeneration time following *stmn2* depletion (Shin et al., 2012). In addition to our findings of delayed distal motor axon degeneration following injury, we observed an impaired ability to regrow motor axons over a 24 hour period which contrast wild type motor axons that displayed a near complete ability to regrow motor axons. Recently it was shown that Stmn2 is essential for motor axons to re-innervate NMJs following sciatic nerve crush in the mouse (Beccari et al., 2025). The findings made in our study support the notion that Stmn2 is required for axon regrowth following injury.

Recent advances in our understanding of misregulated genes in the neurodegenerative disorder amyotrophic lateral sclerosis (ALS) has brought increased attention to the role of STMN2 in disease. In motor neurons from individuals who have succumb to ALS, the RNA binding protein TDP-43 is mislocalized from the nucleus to the cytoplasm where it can form insoluble aggregates (Neumann et al., 2006). TDP-43 (encoded by *TARDBP*) is a ubiquitously expressed members of the heterogeneous nuclear ribonucleoprotein family of proteins that is involved in several steps of RNA processing including of the *STMN2* transcript. In healthy motor neurons, the full-length *STMN2* transcript is composed of 5 exons but following depletion of the RNA binding protein TDP-43, or expression of ALS-associated variants of TDP-43, a cryptic exon in the first intron of *STMN2* termed exon 2a is incorporated in the pre-mRNA that includes a premature polyadenylation site as well as TDP-43 binding motifs (Klim et al., 2019; Melamed et al., 2019; Baughn et al., 2023). This aberrant splicing of the *STMN2* transcript results in the expression of a severely truncated and non-functional protein and reduce expression of full length STMN2 (Baughn et al., 2023). While zebrafish and murine genomes do not possess the corresponding cryptic *STMN2* exon 2a, these comparative models can serve as useful tools for investigating the consequences that loss of *stmn2* expression has on nervous system function *in vivo*. In summary, our findings demonstrate that loss of *stmn2a* and *stmn2b* expression results in a minor motor phenotype in zebrafish larvae and impairment in the ability to regenerate motor axons following injury.

